# An *in vitro* System for Studying Osteochondrogenic Differentiation of Smooth Muscle Cells and Modeling Intimal Vascular Calcification

**DOI:** 10.64898/2026.09.05.749615

**Authors:** João P. Monteiro, Matthew D. Worssam, Wenduo Gu, Daniel Y. Li, Quanyi Zhao, Alex K. Karius, Markus Ramste, Guyu Tracy Qin, Isabella Damiani, Siwen Zheng, Trieu Nguyen, Brian T. Palmisano, Chad S. Weldy, Juyong Brian Kim, Minna U. Kaikkonen, Paul Cheng, Thomas Quertermous

**Author notes:** These authors contributed equally to this manuscript (João P. Monteiro, Matthew D. Worssam). Corresponding Author: Thomas Quertermous, Falk Cardiovascular Research Building CV 297, 870 Quarry Road, Stanford, CA 94305.

## Abstract

**Objective:** Smooth muscle cells (SMCs) undergo phenotypic transitions during atherosclerosis, including towards a chondromyocyte (CMC) state associated with intimal calcification. Although standard *in vitro* calcification assays robustly reproduce mineral deposition, it remains unclear how well they recapitulate these disease-associated SMC states. We sought to define the CMC transcriptional phenotype in atherosclerosis and develop an *in vitro* system that faithfully reproduces it.

**Approach and Results:** We firstly identified a CMC transcriptional signature in murine and human atherosclerotic plaque through single-cell RNA-sequencing, and spatial transcriptomics. CMCs showed a conserved osteochondrogenic program which localized within plaques and adjacent to calcified regions.

We then developed an osteochondrogenic differentiation (OCD) assay by combining well-established calcification components with a high-density SMC micromass culture and TGF-β1 supplementation and benchmarked it against a standard calcification (SC) assay using calcium quantification and bulk RNA-sequencing. Despite comparable calcification, OCD and SC resulted in distinct transcriptional states, with OCD showing preferential upregulation of osteochondrogenic programs, and a higher CMC signature score. Additionally, OCD upregulated genes with a stronger enrichment near coronary artery disease (CAD)-associated loci. These responses were reproducible across several primary human SMC lines. Timecourse analysis also showed that chondrogenic programs preceded calcification and showed directional concordance with the inferred *in vivo* SMC-to-CMC trajectory.

To interrogate regulatory pathways controlling this process, we overexpressed the chondrogenic regulator SOX9, which enhanced cartilage and extracellular matrix programs while repressing inflammatory pathways.

Finally, we examined 552 CAD-associated genes nominated across five genome-wide association studies. Of these, 240 were differentially expressed by day 12, and included established SMC regulators as well as a number of candidates not previously characterized in osteochondrogenic SMC transition.

**Conclusions:** The OCD assay results in a strong calcification phenotype together with a disease-associated CMC-like transcriptional state, providing a reliable *in vitro* model for mechanistic investigation of SMC phenotypic transition and prioritization of candidate regulators.

## Introduction

Intimal calcification (IC) is a defining feature of advanced atherosclerotic lesions and acts as a strong predictor of major cardiovascular event risk, independently of traditional risk factors^1^. In addition to passive mineral nucleation associated with apoptosis and extracellular vesicle release, IC is also actively regulated through cell-mediated processes that recapitulate the molecular programs of osteochondrogenic development^2,3^. Early lineage-tracing studies in mice established that lesional cells expressing osteochondrogenic markers are predominantly smooth muscle cell (SMC)-derived^2,4^ and more recently, related SMC-derived osteochondrogenic states have also been identified in human atherosclerotic lesions^5,6^. Moreover, genome-wide association studies (GWASs) have linked coronary artery calcification risk to numerous genes and pathways regulating SMC phenotype^7^, further supporting a conserved role for SMC phenotypic transition in IC.

Building on these observations, lineage-tracing and single-cell transcriptomic analyses across multiple timepoints have provided increasingly detailed resolution of SMC phenotypic transitions during atherosclerosis, identifying a progression from contractile SMCs through an intermediate fibromyocyte (FMC) phenotype toward a chondromyocyte (CMC) state^8^. CMCs preferentially localize to calcified regions of advanced plaques and are characterized by expression of osteochondrogenic regulators such as *Sox9* and *Runx2*, together with cartilage- and calcification-associated genes including *Comp* and *Ibsp*^9,10^. Notably, the FMC-to-CMC transition has been shown to harbor substantial coronary artery disease genetic risk^10^, emphasizing the need for an *in vitro* model that faithfully recapitulates this disease-relevant transition.

Conventional SMC calcification assays usually include supraphysiologic concentrations of inorganic phosphate or β-glycerophosphate, often combined with calcium chloride and osteogenic supplements such as dexamethasone and ascorbic acid to drive matrix maturation and mineralization^11–14^. These systems have served as useful models for investigating the mechanisms that influence vascular calcification and osteogenic gene expression. However, there is considerable heterogeneity in reagent combinations, dosing, duration and culture formats used across different studies, which limits reproducibility and cross-study comparability. Furthermore, these assays are generally developed and assessed based on calcium deposition and the expression of a small number of osteogenic markers, leaving unclear the extent to which the induced SMC phenotypes correspond to disease-associated cell states identified *in vivo*.

The need for more biologically relevant *in vitro* systems is underscored by the recognition that SMC OCD *in vivo* activates a suite of chondrogenic genes typically absent from standard *in vitro* calcification assays^10^. Moreover, it is shaped not only by mineral and metabolite availability, but also by cell fate-regulating signals and 3D microenvironmental cues. These include TGF-β, BMP, and Wnt signaling, along with cell– cell and cell–matrix interactions, which coordinate activation of lineage-defining transcription factors such as SOX9 and RUNX2^15–18^. Among these pathways, canonical TGF-β signaling through SMAD2/3 is a central regulator of chondrogenic differentiation and cartilage extracellular matrix (ECM) production^19,20^ and TGF-β1 has also been shown to induce expression of chondrogenic regulators in vascular SMCs^21^. An *in vitro* system that models these regulatory circuits would provide a critical bridge between high-resolution single-cell atlases and mechanistic functional studies.

Here, we characterize an *in vitro* OCD assay that combines components for osteogenic mineralization with chondrogenic TGF-β1 stimulation in a high-density micromass culture format^19^, and assess its ability to reproduce the molecular features of CMCs identified in the murine and human atherosclerotic plaque. We benchmark this approach against a standard calcification (SC) assay and show that, despite comparable levels of calcification, OCD induces a broader osteochondrogenic phenotype that more closely resembles the transcriptional signature of human CMCs. Importantly, the system is reproducible across multiple human coronary artery SMC lines and is amenable to functional interrogation of regulators of OCD and IC.

## Methods

### Histological Staining and Imaging

To detect calcification, sections were stained with Alizarin Red solution (EMD Millipore, Cat#2003999), washed with deionized water, and mounted using EcoMount (Biocare Medical, Cat#EM897L). For RNAscope, sections were processed using the RNAscope 2.5 HD Detection RED Kit (Advanced Cell Diagnostics, Cat#322360) following the manufacturer’s instructions and mounted using EcoMount. Sections were imaged using an Echo Revolve microscope (Discover Echo) or a BZ-X810 microscope (Keyence).

### Human Coronary Artery SMC Tissue Culture

Primary human coronary artery SMCs (HCASMCs) and hTERT-HCASMCs^22^ were maintained in SmBM supplemented with SmGM-2 SingleQuots (Lonza, Cat#CC-3182). Primary HCASMC lines were purchased from Cell Applications (Cat#350-05a) and used at passages 3-8.

### Induction of SMC Differentiation

For the OCD assay, cells were seeded in a high-density micromass format. Following trypsinization and counting, cells were resuspended in complete media at 5,000 cells/µL, and 10-µL droplets were placed in the center of each well of a 24-well plate. After 2 hours of incubation to allow for cell compaction, 500 µL of complete medium was added to each well. For the SC assay, 75,000 cells were seeded per well in 24-well plates at a density of 150,000 cells/mL. Treatment media were applied the following day.

Basal medium consisted of DMEM containing glucose, L-glutamine, and sodium pyruvate (Corning, Cat#10-013-CV), supplemented with 5% FBS and 100 nM insulin from the SmGM-2 SingleQuot Kit. Phosphate-containing basal medium was prepared by adding β-glycerophosphate (Sigma-Aldrich, Cat#G9422-10G) to a final concentration of 5 mM, followed by sterile filtration.

SC medium was prepared fresh and consisted of phosphate-containing basal medium supplemented with L-ascorbic acid (50 µg/mL; Thermo Fisher Scientific, Cat#036237.14), dexamethasone (10 nM; Sigma-Aldrich, Cat#D2915), and CaCl_2_ (2.5 mM; Thermo Fisher Scientific, Cat#J63122.AE). OCD medium contained all components of the SC medium with the addition of recombinant human TGF-β1 (10 ng/mL; Thermo Fisher Scientific, Cat#100-21-50UG). Media were replaced every 3 days, and cells were maintained under treatment conditions for 3-12 days.

### RNA Extraction and Sequencing

RNA was isolated using the RNeasy Plus Micro Kit (Qiagen, Cat#74034) according to the manufacturer’s instructions. For bulk RNA-sequencing, RNA quality control, library preparation, and sequencing were performed by Novogene. Libraries were sequenced on a NovaSeq X Plus platform to a target depth of 30 million reads/sample.

For qPCR, cDNA was synthesized using the High-Capacity RNA-to-cDNA Kit (Thermo Fisher Scientific, Cat#4388950). qPCR was performed using TaqMan Universal Master Mix II (Thermo Fisher Scientific, Cat#4440040) on a ViiA 7 Real-Time PCR System (Life Technologies).

### Bulk RNA-seq Data Analysis

All raw FASTQ files were evaluated for sequencing quality using *FastQC*. Adapter sequences and low-quality bases were trimmed using *Cutadapt*. Trimmed reads were then aligned to the human reference genome (GRCh38) using *STAR* with the default parameters. Gene-level read counts were generated from the aligned reads using *featureCounts*.

Gene-level differential expression (DE) analysis was performed in R using *DESeq2*. Genes with a log2-fold-change >0.5 and adjusted *P*-value <0.05 were considered significantly differentially expressed. For comparisons between SC and OCD, each treatment condition was compared with its corresponding control condition. For the OCD time course, samples collected at days 3, 6, and 12 were compared with their respective time-matched controls. For visualization and unsupervised analysis presented, count data was variance-stabilizing transformed using *DESeq2*. Principle component analysis was performed on variance-stabilized expression values to evaluate global transcriptional differences between experimental conditions. Volcano plots were generated using *EnhancedVolcano* from *DESeq2* log2-fold-change and adjusted *P-values* DE results. Heatmaps were generated from variance-stabilized expression values. For analyses of transcriptional patterns across experimental conditions and the OCD time course, the 500 most variable genes were selected based on variance across samples and grouped into four clusters by k-means clustering. Expression values were scaled by gene for visualization. Transcription factor activity was inferred using human *DoRothEA* regulons (confidence levels A–C) and the univariate linear model method in *decoupleR* (run_ulm; minimum regulon size, 5), using *DESeq2* Wald test statistics as input. For GWAS enrichment analysis, variants were obtained from the NHGRI-EBI GWAS Catalog and consolidated overlaping traits into 9 disease categories. Differentially expressed genes were converted to intervals spanning 5 kb upstream to 2kb downstream of the TSS and intersected with category variants using BEDTools. Fold enrichment over the genome-wide expectation was calculated as previously described^5^.

### Consensus Non-Negative Matrix Factorization Analysis

Consensus non-negative matrix factorization (cNMF) was performed as previously described^23,24^ using a human arterial single-cell RNA sequencing (scRNA-seq) dataset, comprising 9 distinct arterial sites from 6 patients^5^. The top 100 genes from the CMC-associated program were used to define the CMC signature for downstream analyses.

### Xenium slide processing and analysis

Human coronary artery sections were profiled using the 10x Genomics Xenium Prime 5K gene expression panel. Sample processing and downstream analysis were performed as previously described^10^. The CMC transcriptional signature was quantified as a module score across spatially resolved cells. Cells with a module score ≥1.3 were classified as CMC-signature positive.

### Calcium Quantification

At the end of differentiation, culture media were removed and wells were washed with PBS. Freshly prepared 0.6 N HCl was added to each well to solubilize calcium deposits, and supernatant was collected after 48 hours of incubation at 4°C. Wells were then washed once with PBS, and residual cellular material was lysed with 0.1 N NaOH containing 0.1% SDS for 30 minutes at room temperature for total protein extraction.

Calcium concentration in the acid extracts was measured using the QuantiChrom Calcium Assay Kit (BioAssay Systems, Cat#DICA-500). Total protein content in the NaOH/SDS lysates was measured using the Pierce BCA Protein Assay Kit (Thermo Fisher Scientific, Cat#23225). Calcium content was normalized to total protein for each sample.

For visualization of calcification, cells were fixed with 4% PFA for 10 minutes and washed three times with PBS for 5 minutes each. Cells were stained with Alizarin Red solution for 20 minutes at room temperature on a rocker, followed by three 5-minute washes with deionized water. Wells were imaged using a Keyence microscope.

### Data availability

Murine atherosclerotic plaque single-cell RNA-sequencing data were obtained from Gene Expression Omnibus (GEO) accession GSE321762^10^, and human arterial single-cell RNA-sequencing data were obtained from CELLxGENE: 8f17ac63-aaba-44b5-9b78-60f121da4c2f^5^. Murine aorta spatial transcriptomic Xenium data were obtained from GEO accession GSE316666^10^.

## Results

### Smooth Muscle Cell Phenotypic Transition Toward a Chondromyocyte Fate Is Defined by a Conserved Transcriptional Program

To establish a disease-relevant molecular benchmark for evaluating *in vitro* models of SMC OCD, we first examined the transcriptional features of CMCs in murine and human atherosclerotic plaques. Our scRNA-seq analysis resolved SMCs into distinct contractile SMC, FMC, and CMC populations in both species (Figure 1Ai). CMCs were distinguished from contractile SMCs and FMCs by increased expression of osteochondrogenic regulators and ECM genes, including *SOX9, COMP, RUNX2*, and *IBSP*. To more comprehensively define the CMC transcriptional state, we applied cNMF to a scRNA-seq dataset of human plaque from 6 patients and identified a 100-gene transcriptional program associated with the CMC population^23,24^. Module scoring of this CMC signature showed significant and comparable enrichment within the CMC populations of both human and murine lesions, supporting conservation of this transcriptional state across species (Figure 1Ai).

**Figure 1.**
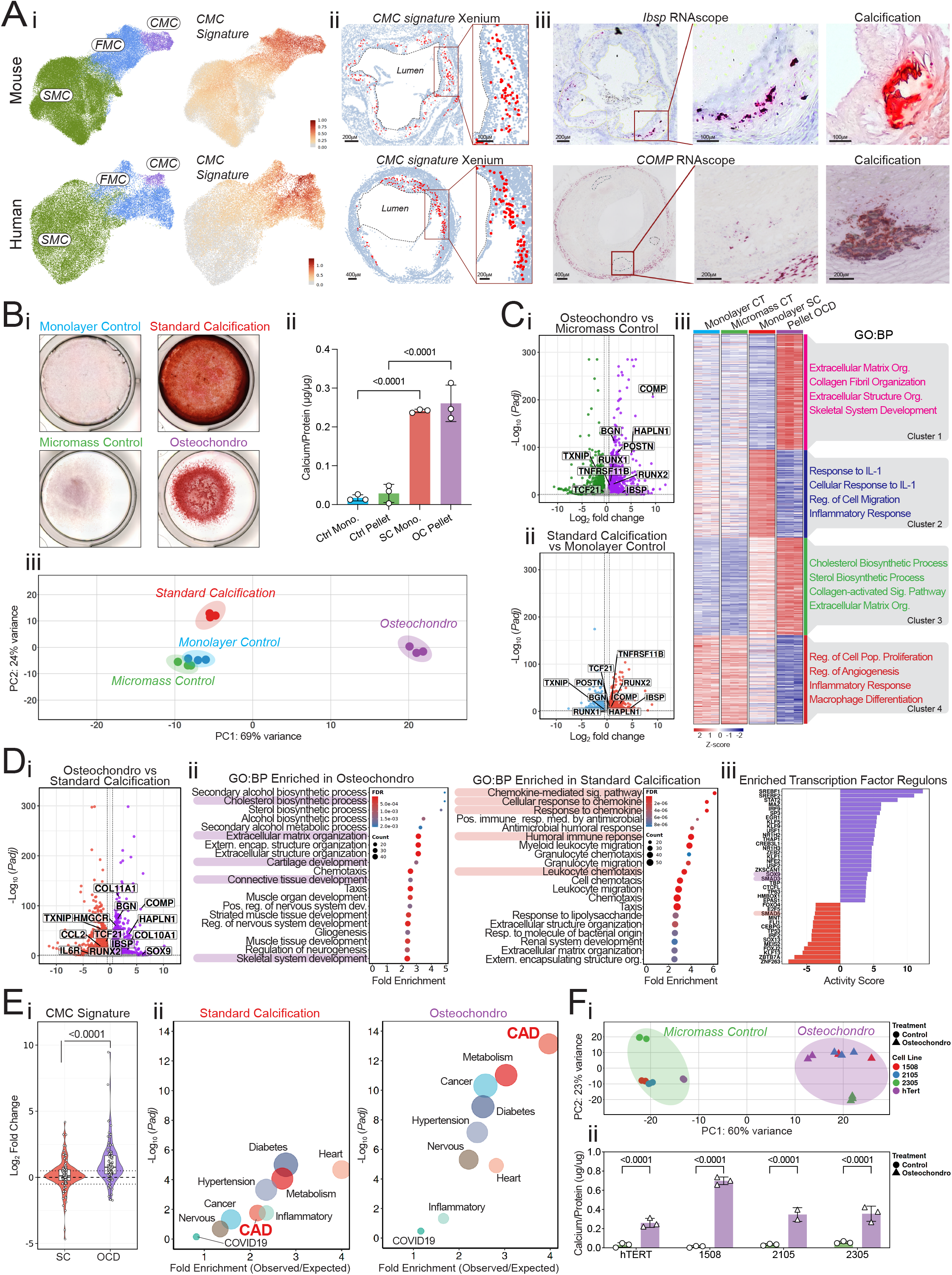
Osteochondrogenic differentiation assay recapitulates the transcriptional identity of chondromyocytes observed in atherosclerotic plaque. **(A) Conserved gene expression in murine and human atherosclerotic plaque. (i)** Uniform manifold approximation and projection (UMAP) plots of single-cell RNA sequencing datasets from murine (top) and human (bottom) atherosclerotic plaque identify contractile smooth muscle cells (SMCs), fibromyocytes (FMCs), and chondromyocytes (CMCs). Feature plots show module score for human CMC gene cNMF program and *COMP* expression across the murine (top) and human (bottom) SMC UMAPs. **(ii)** 10x Genomics Xenium images of 16-week HFD murine aorta (top) and human coronary artery (bottom) showing cells positive for the human CMC transcriptional signature. **(iii)** Representative sequential sections from murine (top right) and human (bottom right) atherosclerotic lesions were analyzed by RNAscope for *COMP* and *IBSP* and by Alizarin Red staining for calcification. **(B) Assessment of calcification induced by SC and OCD. (i)** Representative Alizarin Red-stained samples at day (D) 12 are shown for control monolayer, SC monolayer, control micromass, and OCD micromass conditions. **(ii)** Calcium deposition was quantified by acid extraction and normalized to total protein content. Data are presented as mean ± SEM (n=3). Statistical significance determined by one-way ANOVA followed by Bonferroni’s multiple comparisons test. **(iii)** Principal component analysis (PCA) of bulk RNA sequencing data from control monolayer, standard calcification (SC), control micromass, and osteochondrogenic differentiation (OCD) conditions at D12 (n=3). **(C) Transcriptomic comparison of OCD and SC with their respective controls at day 12. (i)** Volcano plots show differential expression (DE) for OCD and **(ii)** SC relative to respective controls. **(iii)** The heatmap displays the 500 most variable genes across all four conditions following variance-stabilizing transformation and k-means clustering into four clusters. Gene Ontology Biological Process (GO:BP) enrichment analysis is shown for each gene cluster. **(D) Direct transcriptomic comparison of OCD and SC at D12. (i)** Volcano plot shows DE genes in OCD versus SC. **(ii)** GO:BP enrichment analysis is shown separately for genes expressed more highly in OCD and genes expressed more highly in SC. **(iii)** DoRothEA regulon analysis shows inferred transcription factor activity in OCD relative to SC (right), including *SOX9* and *RUNX2*. **(E) Human CMC gene program and GWAS enrichment analysis of OCD and SC at D12. (i)** Violin plots show log_2_ fold-change of genes in the human CMC cNMF program for SC and OCD relative to respective controls. Data are presented as log_2_ fold-change values for individual, with boxplots indicating median and interquartile range. Statistical significance was determined by paired Wilcoxon signed-rank test. **(ii)** Genome-wide association study (GWAS) enrichment analysis of genes upregulated in SC and OCD relative to respective controls, across 9 disease categories. **(F) Calcification and transcriptional responses across human SMC lines. (i)** PCA of bulk RNA-sequencing data from three independent primary human coronary artery smooth muscle cell (HCASMC) donors and immortalized hTERT-HCASMCs following OCD. **(ii)** Normalized calcification produced by each HCASMC line at D12 of OCD and control media treatment. Data are presented as mean ± SEM (n=3). Statistical significance determined by one-way ANOVA followed by Bonferroni’s multiple comparisons test.

Spatial transcriptomic profiling further demonstrated that cells enriched for the CMC signature were concentrated within the plaque core for both murine and human atherosclerotic lesions, with little enrichment in the media or fibrous cap (Figure 1Aii). Consistent with this, we observed expression of CMC-associated genes *COMP* and *IBSP* adjacent to calcified regions at the base of murine and human lesions, highlighting the association between this transcriptional state and IC (Figure 1Aiii).

Together, these data define a conserved and spatially restricted CMC-associated transcriptional program in atherosclerosis and provide a molecular reference against which *in vitro* models of SMC OCD and calcification can be evaluated.

### The Osteochondrogenic Differentiation Assay Induces Calcification and Recapitulates CMC-Associated Transcriptional Programs

To promote osteochondrogenic SMC differentiation, we developed an OCD assay that combines calcification-promoting components with chondrogenic high-density micromass cell culture format and TGF-β1 supplementation. To benchmark this model, we compared it with a standard calcification (SC) assay. We first assessed mineralization by Alizarin Red staining after 12 days of differentiation and observed robust and comparable calcification with both assays (Figure 1Bi-ii). To determine whether the OCD assay recapitulates *in vivo* CMC transcriptional changes and how this response compares with SC assay conditions, we performed RNA-seq on humanized-TERT (hTERT)-HCASMCs after 12 days of OCD/SC treatment, alongside their respective controls. Principal component analysis (PCA) showed consistency between experimental replicates, but there was a clear separation between assays, with OCD samples forming a distinct transcriptional cluster from both control and SC groups (Figure 1Biii).

Further, differential expression analysis comparing OCD and SC treatments with their respective controls demonstrated substantial transcriptional reprogramming under both conditions (Figure 1Ci-ii). While SC and OCD shared upregulation of several classical osteogenic genes, including *RUNX2, IBSP*, and *COL10A1*, the magnitude of their upregulation was markedly higher under OCD conditions. Moreover, OCD activated a broader osteochondrogenic program, characterized by robust induction of cartilage matrix and chondrocyte-associated genes, including *HAPLN1, MATN3, COL11A1, MMP13*, and *NKX3-2*, which was absent from the SC assay. (Figure 1Ci-ii).

Analysis of the 500 most variable genes across all four conditions highlighted distinct treatment-associated expression patterns segregated into four k-means clusters (Figure 1Ciii). The cluster most strongly associated with OCD included osteochondrogenic genes *COMP, COL10A1, COL11A1, HAPLN1, MATN3, CILP*, and *CILP2*, with Gene Ontology Biological Process (GO:BP) term enrichment for ECM organization, and skeletal system development. In contrast, clusters more prominent under SC included inflammatory genes such as *CHI3L1, IL1R1, CCL2*, and *CXCL1*, with concordant GO:BP term enrichment. (Figure 1Ciii).

To define the transcriptional features that distinguish the two induced states, we also directly compared OCD and SC at day 12. Genes expressed more highly in OCD were enriched for ECM organization, cartilage, connective tissue and skeletal system development programs, whereas genes expressed more highly under SC were enriched for inflammatory processes, including chemokine signaling, and leukocyte chemotaxis (Figure 1Di-ii). Thus, the two assays generate transcriptionally distinct SMC phenotypes despite achieving similar calcification.

Transcription factor activity analysis additionally distinguished between these phenotypes. Regulon analysis identified increased SOX9 activity in OCD relative to SC, consistent with its role as a chondrogenic master regulator^25^ (Figure 1Diii). Importantly, RUNX2 did not emerge as differentially active between conditions, as its activity represented a common osteogenic feature of both OCD and SC. We also saw increased SMAD3/4 regulon activity in OCD, while SMAD5 activity was decreased consistent with canonical TGF-β1-driven SMAD2/3 signaling introduced by the OCD treatment, rather than a BMP-dominant SMAD1/5/8 response^17,26,18^.

To determine whether the OCD assay reproduces the transcriptional identity of CMCs observed *in vivo*, for each gene within the human CMC signature, we calculated the fold change for OCD/SC-treated cells relative to their respective controls. CMC signature genes showed significantly higher induction under OCD than SC, with a mean log_2_ fold-change of 1.08 versus 0.15 (Figure 1Ei), thus indicating substantially greater transcriptional similarity between OCD-treated SMCs and human CMCs.

Finally, we compared enrichment of OCD- and SC-upregulated genes across GWAS loci grouped into 9 disease categories. For OCD, CAD loci showed the highest enrichment of any category (3.97 fold), whereas for SC, CAD ranked seventh of ten (2.16 fold), behind cardiomyophaty, diabetes and metabolism traits (Figure 1Eii). This preferential enrichment further supports the ability of the OCD assay to capture gene expression changes associated with SMC phenotypic transition in human atherosclerosis.

### The Osteochondrogenic Differentiation Assay Produces Reproducible Calcification and Transcriptional Responses Across Human SMC Lines

To assess the robustness of the OCD assay across genetically distinct human backgrounds, we applied the same protocol to primary HCASMC lines from three independent donors, together with the immortalized hTERT-HCASMC line used previously. Despite donor-specific differences in baseline transcriptional state, OCD produced a consistent shift in global gene expression across the independently derived HCASMC lines, including potent upregulation of human CMC signature genes *COMP, HAPLN1* and *IBSP*. (Figure 1Fi-ii).

### Timecourse Analysis Shows Ordered Molecular Events During Osteochondrogenic SMC Transition

To characterize the progression of OCD, we measured calcification and performed transcriptomic profiling at multiple differentiation timepoints. An early osteochondrogenic and ECM response was evident at day 3 (D3), with induction of *COMP, CILP2, MATN3, COL10A1, CILP, HAPLN1*, and *MMP13*, together with more modest induction of the osteogenic transcription factor *RUNX2* (Figure 2Ai). This response became more pronounced at later timepoints, with increased expression of *COMP* and *COL10A1* and expansion of the program to include genes associated with chondrocyte maturation and calcification, including *COL11A1, NKX3-2, PTHLH, ANKH, ENPP1, PHOSPHO1*, and *BMPR1B*. By D12, expression of chondrogenic genes *COMP* and *COL10A1* reached >1000-fold increase relative to control conditions. In parallel, the contractile SMC regulator *MYOCD* and FMC-associated genes, including *TCF21* and *TXNIP*, were reduced during differentiation. Inflammatory genes were also suppressed during OCD, particularly at D6, replicating the reduction of inflammatory programs during FMC-to-CMC transition in atherosclerotic plaque (Figure 2Ai).

**Figure 2.**
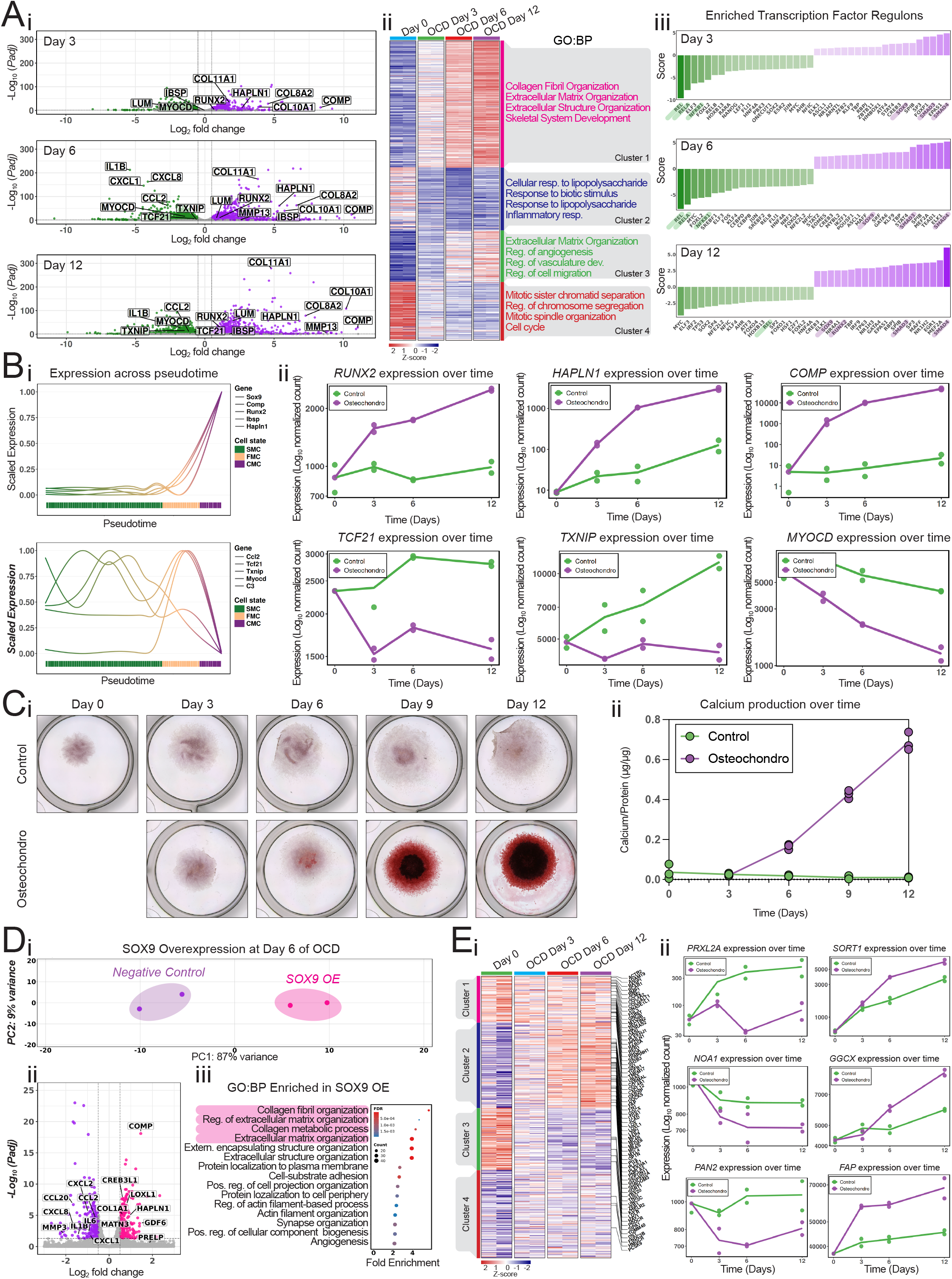
Osteochondrogenic differentiation assay reveals ordered transcriptional dynamics and enables functional interrogation of SMC osteochondrogenesis. **(A) Transcriptomic profiling across the osteochondrogenic differentiation (OCD) timecourse. (i)** Volcano plots show differential expression (DE) at days 3, 6, and 12 relative to time-matched control conditions. **(ii)** The heatmap displays the 500 most variable genes across the OCD time course grouped into four clusters by k-means clustering (middle), with corresponding Gene Ontology Biological Process (GO:BP) enrichment analysis. **(iii)** DoRothEA analysis shows inferred activity of 40 transcription factors at each OCD time point relative to the corresponding time-matched control (right). **(B) Comparison of gene-expression patterns across *in vivo* SMC pseudotime and the *in vitro* OCD time course. (i)** Scaled gene expression across pseudotime is shown for lineage-traced SMC-derived cells collected across seven high-fat-diet time points (left). Genes shown include osteochondrogenic-associated genes (*Sox9, Runx2, Comp, Ibsp, Hapln1, Snai1, Runx1*, and *Col10a1*), contractile SMC regulator gene (*Myocd*), and FMC- or inflammatory-associated genes (*Tcf21, Txnip, Ccl2, Ccl7, Cxcl12, Fbln1*, and *C3*). Each curve shows a generalized additive model fit to log-normalized expression across all cells and min–max scaled to the interval 0–1, showing relative rather than absolute expression. Pseudotime is in arbitrary units, reflecting each cell’s inferred position along the transition. Line color indicates position along pseudotime, transitioning from green (SMC) through orange (FMC) to purple (CMC). The horizontal band beneath the axis indicates the most frequent cell state at each point along pseudotime. **(ii)** Expression across four OCD timepoints is shown for selected osteochondrogenic-associated genes (*RUNX2, HAPLN1*, and *COMP*), FMC-associated genes (*TCF21* and *TXNIP*), and the contractile SMC regulator *MYOCD* (right). **(C) Calcification across the OCD timecourse. (i)** Representative Alizarin Red-stained hTERT-HCASMC micromasses are shown at days 0, 3, 6, and 12 of control or OCD-media treatment. **(ii)** Calcium deposition was quantified at the same four time points by acid extraction and normalized to total protein content. **(D) *SOX9* overexpression during OCD. (i)** PCA of bulk RNA-sequencing data from hTERT-HCASMCs overexpressing the chondrogenic regulator *SOX9* or a negative lentiviral control expressing a size-matched *Luciferase* construct during OCD at day 6. **(ii)** Volcano plot shows DE genes in *SOX9*-overexpressing cells relative to negative-control cells. **(iii)** GO enrichment analysis is shown for genes upregulated following SOX9 overexpression. **(E) Expression of coronary artery disease (CAD)-associated genes during OCD. (i)** Heatmap shows expression of 552 CAD-associated genes nominated in at least one of five genome-wide association studies^27–31^ across OCD assay days 0, 3, 6, and 12; 240 of these genes were significantly DE at day 12 relative to time-matched controls. **(ii)** Expression across four OCD timepoints is shown for selected CAD-associated genes that were upregulated (*SORT1, GGCX*, and *FAP*) or downregulated (*PRXL2A, NOA1*, and *PAN2*) during OCD.

K-means clustering grouped the 500 most variable genes into four clusters with distinct temporal expression patterns (Figure 2Aii). Consistent with a progressive establishment of the osteochondrogenic phenotype described above, GO:BP analysis showed that Cluster 1 was enriched for ECM organization, and skeletal system development. In contrast, Cluster 2, containing genes suppressed during OCD, was enriched for inflammatory response, while the remaining clusters were associated with vascular remodeling and cell-cycle processes (Figure 2Aii). Regulon analysis further identified coordinated changes in inferred transcription factor activity during OCD (Figure 2Aiii). SMAD3, SMAD4, and SOX9 showed increased inferred activity at all timepoints, indicating early and sustained activation of chondrogenic regulatory programs. RUNX2 activity was additionally increased at D12, consistent with the emergence of a stronger osteogenic program at the later stage of differentiation. In contrast, NF-κB family members, including RELA, REL, and NFKB1, showed strongly reduced activity at earlier timepoints, consistent with the early suppression of inflammatory gene programs observed in the differential expression and GO analyses.

To determine how these *in vitro* transcriptional changes related to SMC phenotypic modulation during atherosclerosis, we compared the OCD timecourse with an *in vivo* pseudotime trajectory of lineage-traced SMC-derived cells collected across seven timepoints in a murine model of atherosclerosis^10^ (Figure 2Bi).

Along the *in vivo* trajectory from contractile SMC through FMC and toward CMC states, osteochondrogenic-associated genes including *Sox9, Runx2* and *Comp* increased as cells approached CMC identity. In contrast, the contractile regulator *Myocd* and FMC- or inflammatory-associated genes, including *Tcf21, Ccl2*, and *C3*, declined or showed maximal expression in intermediate states. Consistent with the *in vivo* dynamics, in the OCD timecourse, *RUNX2, HAPLN1*, and *COMP* were expressed more highly at later time points, whereas *TCF21, TXNIP*, and *MYOCD* declined or remained low throughout (Figure 2Bii). Overall, the concordance of temporal gene expression patterns between OCD and atherosclerosis indicates that OCD recapitulates disease-relevant SMC-derived cell states.

Lastly, in concert with increasingly upregulated osteochondrogenic transcriptional programs, calcification developed progressively, with minimal mineralization at D0-3, but steadily increasing by D6-D12 (Figure 2Ci-ii).

Collectively, these results indicate that the OCD assay captures an ordered phenotypic transition marked by progressive osteochondrogenic transcriptional changes, culminating in robust calcification.

### Functional Interrogation of SMC Phenotypic Transition Using the Osteochondrogenic Differentiation Assay

To determine whether the OCD assay could detect transcriptional consequences of experimental manipulation, we overexpressed the central chondrogenic regulator *SOX9* in hTERT-HCASMCs and compared with a negative lentiviral control during OCD. At D6, PCA demonstrated clear separation of *SOX9*-overexpressing cells from control treated cells, indicating a broad effect of *SOX9* on the transcriptional state acquired during OCD, consistent with the previously observed activation of the SOX9 regulon during OCD (Figure 1Diii, 2Di).

DE analysis showed increased expression of cartilage- and ECM-associated genes following *SOX9* overexpression, including *COMP, GDF6, HAPLN1*, and *MATN3*, together with matrix-remodeling genes such as *ADAMTSL2*, and *THBS2*. Notably, *RUNX2* and *IBSP* were not among the DE genes, suggesting that SOX9 preferentially reinforced the chondrogenic and matrix-producing components of the OCD program rather than broadly increasing osteogenic differentiation. In parallel, inflammatory genes including *IL1B, CXCL8, CCL7*, and *CCL2* were downregulated. Consistent with these gene-level changes, GO:BP analysis showed enrichment of ECM and collagen organization programs together with suppression of inflammatory pathways (Figure 2Dii-iii).

The coordinated transcriptional response to manipulation of an established chondrogenic regulator demonstrates that the OCD assay can be used not only to characterize SMC phenotypic transition, but also to functionally interrogate regulatory pathways controlling this process.

### Osteochondrogenic Differentiation as a High-Throughput Tool for CAD GWAS Prioritization

To assess the utility of the OCD assay for identifying disease-relevant regulators of SMC phenotypic transition, we examined the expression of 552 CAD-associated genes compiled from five large-scale GWASs^27–31^. These genes exhibited heterogeneous temporal expression patterns across OCD, with 240 genes significantly differentially expressed in OCD samples at D12 relative to time-matched controls (Figure 2Ei). These included established causal CAD genes with known roles in SMC phenotypic modulation, such as *TCF21, KLF4*, and *PLPP3*, supporting the ability of the assay to capture disease-relevant SMC biology.

The analysis also highlighted genes whose roles in SMC OCD remain incompletely characterized, including genes upregulated during the assay, such as *SORT1, GGCX*, and *FAP*, and downregulated genes, including *PRXL2A, NOA1*, and *PAN2* (Figure 2Eii). The regulation of these genes during acquisition of the

CMC-like transcriptional state nominates them as candidate mediators or markers of SMC phenotypic transition and provides a basis for subsequent functional investigation.

## Discussion

SMCs undergo complex phenotypic transitions during atherosclerosis that localize to discrete regions of the plaque and are proposed to exert distinct effects on disease progression and risk^10,32,33^. Although *in vivo* single-cell studies have defined transcriptional and epigenomic programs associated with these transitions, mechanistic investigation has been limited by the absence of *in vitro* systems that faithfully recapitulate disease-associated cell states. In the present study, we establish the OCD assay as a model of the osteochondrogenic CMC state associated with atherosclerotic IC and the intermediate transitions leading towards this phenotype. Importantly, we demonstrate that comparable levels of calcification can arise from markedly different transcriptional states. Although both OCD and SC assays induced robust osteogenic gene expression, OCD additionally activated a broader chondrogenic and ECM program, which also induced concomitant transcriptional changes resembling the CMC-associated transcriptional state observed in murine and human atherosclerotic plaque. Thus, calcification alone does not define the molecular phenotype being modeled, and transcriptional identity should be considered when selecting *in vitro* systems to study vascular calcification.

This distinction has important implications for modeling intimal versus medial calcification. CMCs in our *in vivo* datasets localized to calcified regions within the atherosclerotic intima, and genes regulated by OCD showed strong enrichment near CAD-associated GWAS loci, whereas this enrichment was weaker under SC conditions. In contrast, SC produced an osteogenic response accompanied by prominent inflammatory programs and lacked comparable activation of chondrogenic programs. Standard *in vitro* calcification models are widely used to study the osteogenic conversion of SMCs under conditions relevant to CKD-associated medial calcification and activate RUNX2 together with NF-κB-dependent inflammatory signaling^34^. Recent *in vivo* studies of medial calcification similarly demonstrate SMC osteogenic conversion accompanied by inflammatory signaling^35,36^. While direct molecular comparison with medial calcification tissue will be required to establish this distinction formally, these observations suggest that SC and OCD should be viewed as complementary rather than interchangeable models, tailored for medial and atherosclerotic intimal calcification respectively.

Defining the precise sequence of the molecular changes regulating pathogenic SMC transitions *in vivo* is challenging because atherosclerotic lesions contain heterogeneous SMC-derived populations at multiple stages of transition. While single-cell trajectory analyses can infer how these states are ordered, the OCD assay offers a controlled system in which the temporal sequence of these changes can be directly studied. Through the timecourse, the chondrogenic and ECM programs appeared before substantial calcification was present and followed the same directional pattern as that inferred from the *in vivo* SMC trajectory, indicating that the assay may reflect a progressive remodelling toward CMC identity rather than a terminal calcified state. This response was reproducible across independent primary HCASMC backgrounds, supporting its broad application across HCASMC cell lines. In addition, the overlap between genes regulated during OCD and CAD GWAS-associated genes provides a framework for prioritizing candidate regulators of osteochondrogenic SMC transition for subsequent functional and *in vivo* investigation.

Despite its usefulness, the OCD assay is still a reductionist model and is not intended to reproduce the complex environment of atherosclerotic plaque biology. It does not replicate hemodynamic forces, immune-cell interactions, lipid accumulation, hypoxia, or the spatial signaling gradients present within lesions, which influence SMC state transitions *in vivo*. The assay should therefore be viewed as a complementary experimental system for prioritizing and functionally assessing candidate regulators before using more resource-intensive *in vivo* models. Incorporating additional plaque-associated cues, co-culture systems, mechanical stimuli, or single-cell profiling could further increase its physiological relevance and help determine which features of the CMC phenotype can be attributed to specific environmental signals.

In summary, we describe an *in vitro* osteochondrogenic SMC differentiation assay that recapitulates the CMC transcriptional state associated with atherosclerotic intimal calcification. Its principal advantage over conventional calcification assays is not greater calcium deposition, but rather greater transcriptional similarity to atherosclerotic SMC cell transition states and CAD associated gene expression, thus providing a tractable *in vitro* system for mechanistic investigation of SMC phenotypic transition.

## Acknowledgements

This work was supported by the American Heart Association grants 24POST1187860 (JPM), 26CDA1596032 (JPM), 25POST1360338 (MDW), 24CDA1272805 (BTP), National Institutes of Health grants R01HL139478 (TQ), R01HL134817 (TQ), R01HL171045 (TQ), R01HL158525 (TQ), UM1HG011972 (TQ), K08HL177251 (BTP), a Network Grant from the LeDucq Foundation, and the William G. Irwin Foundation (TQ), as well as a Human Cell Atlas grant (ZF2019-002437) from the Chan Zuckerberg Foundation (PC, TQ).

